# BioImage Model Zoo: A Community-Driven Resource for Accessible Deep Learning in BioImage Analysis

**DOI:** 10.1101/2022.06.07.495102

**Authors:** Wei Ouyang, Fynn Beuttenmueller, Estibaliz Gómez-de-Mariscal, Constantin Pape, Tom Burke, Carlos Garcia-López-de-Haro, Craig Russell, Lucía Moya-Sans, Cristina de-la-Torre-Gutiérrez, Deborah Schmidt, Dominik Kutra, Maksim Novikov, Martin Weigert, Uwe Schmidt, Peter Bankhead, Guillaume Jacquemet, Daniel Sage, Ricardo Henriques, Arrate Muñoz-Barrutia, Emma Lundberg, Florian Jug, Anna Kreshuk

## Abstract

Deep learning-based approaches are revolutionizing imaging-driven scientific research. However, the accessibility and reproducibility of deep learning-based workflows for imaging scientists remain far from sufficient. Several tools have recently risen to the challenge of democratizing deep learning by providing user-friendly interfaces to analyze new data with pre-trained or fine-tuned models. Still, few of the existing pre-trained models are interoperable between these tools, critically restricting a model’s overall utility and the possibility of validating and reproducing scientific analyses. Here, we present the BioImage Model Zoo (https://bioimage.io): a community-driven, fully open resource where standardized pre-trained models can be shared, explored, tested, and downloaded for further adaptation or direct deployment in multiple end user-facing tools (e.g., ilastik, deepImageJ, QuPath, StarDist, ImJoy, ZeroCostDL4Mic, CSBDeep). To enable everyone to contribute and consume the Zoo resources, we provide a model standard to enable cross-compatibility, a rich list of example models and practical use-cases, developer tools, documentation, and the accompanying infrastructure for model upload, download and testing. Our contribution aims to lay the groundwork to make deep learning methods for microscopy imaging findable, accessible, interoperable, and reusable (FAIR) across software tools and platforms.

## Main

Since the first major success of modern deep learning (DL) on the ImageNet Large Visual Recognition Challenge, convolutional neural networks (CNNs) have redefined the state-of-the-art on virtually all open computer vision problems. Prominent examples include tasks as diverse as image synthesis^1^, scene understanding^2^, and image enhancement^3^, but the overall effect of DL technology has been so extensive that it is difficult to find an image-related task that has not profoundly benefited from it. Microscopy imaging has been no exception: DL-based methods in image reconstruction^4–7^, classification^8–10^, segmentation^11–13^, and artificial labeling^14,15^ have enabled both flagship projects^16–19^ and common “bread-and-butter” tasks, allowing image analysis to keep pace with recent advancements in imaging technology and instrumentation.

Deep neural networks are controlled by millions of parameters whose values need to be found during the training process. While this parameterization enables trained DNNs to generalize to previously unseen data, it makes the training process very computationally intensive as well as time- and data-consuming. It is therefore common practice to use a network that was previously trained on the desired task and re-adjust it to new data by a few additional training iterations. The value of pre-trained networks is substantial: a great share of newly published methods exploit popular architectures pre-trained on the largest available public datasets^20^. Companies and research groups maintain their collections with publicly shared networks organized into the so-called *model zoos* (https://pytorch.org/hub/, https://www.tensorflow.org/hub, https://modelzoo.co/, https://huggingface.co/). With the growing adoption of AI-based methods, new collections have started to appear for biomedical data, i.e. in genomics (https://kipoi.org/), single-cell transcriptomics (https://sfaira.readthedocs.io/en/latest/), or medical imaging (https://monai.io/).

Nevertheless, the existing model repositories contain very few networks trained on microscopy data. On one hand this is because many repositories only take models trained on publicly available data, on the other hand because most model zoos are designed for method developers, not users (i.e. microscopists). When submitting a model to a model zoo, computer scientists from the natural image domain or bioinformatics primarily address other developers who will use the models in their code and deploy them without requiring any further help. In contrast, a new microscopy image analysis method creates impact by reaching out to end-users: scientists in experimental labs and imaging facilities who benefit from point-and-click tools in combination with scripting^21^. Thus, wide adoption by life scientists will only efficiently occur when DL solutions can be directly called from a convenient graphical user interface (GUI) or, better yet, if they are integrated into existing and familiar image analysis software suites. The integration is, however, far from trivial and needs to be performed time and time again for each independent software tool.

In recent years, a growing community of DL method developers has gone far beyond simply publishing the source code of their new ideas. New Fiji/ImageJ^22,23^ or napari^32^ plugins (CSBDeep^24^, StarDist^25,26^, U-Net^27^) and standalone GUI-based tools (CellPose^12^, NucleAIzer^11^, DeepCell^26^) have pushed both the performance and the accessibility of the DL state-of-the-art for microscopy image restoration and segmentation. ImJoy^29^ was developed to ease the sharing and deployment of deep learning tools by providing a scalable and interactive plugin framework backed by the web and Python ecosystem. DeepImageJ^30^ brought many popular CNNs to the broad user base of Fiji, allowing to seamlessly combine network inference with other analysis steps available within Fiji’s UI. ZeroCostDL4Mic^31^ has democratized the training process, making an extensive collection of networks available through user-friendly Google Colab notebooks and demonstrating the power of cloud computing infrastructure for quick network evaluation. While such tools provide tremendous value to life science researchers, their siloed network collections are confined to execution in individual tools, severely limiting the reusability, reproducibility, and interoperability of DL models in microscopy image analysis pipelines.

To address the limitations listed above, we present the BioImage Model Zoo (Figure 1), a model repository tailored to the needs of the whole microscopy image analysis community. At the core of our developments lies a unified way of describing and consuming trained DL models. This is achieved through a standard model description format that captures all necessary model metadata, including input and output data format, pre-trained weight values and training data provenance. Our libraries allow for standardized model execution with minimal code and easy integration into user-facing tools and frameworks. The BioImage Model Zoo is already supported by ilastik^33^, deepImageJ^22^, ImJoy^29^, StarDist^16,17^, ZeroCostDL4Mic^31^, and QuPath^34^. We are now ready to welcome new community partners: our ambition is to make the uptake of our metadata formats so straightforward that any method developer or coding user will be able to contribute models to the Zoo and enrich their own tools by consuming the models from our repository.

**Figure 1.**
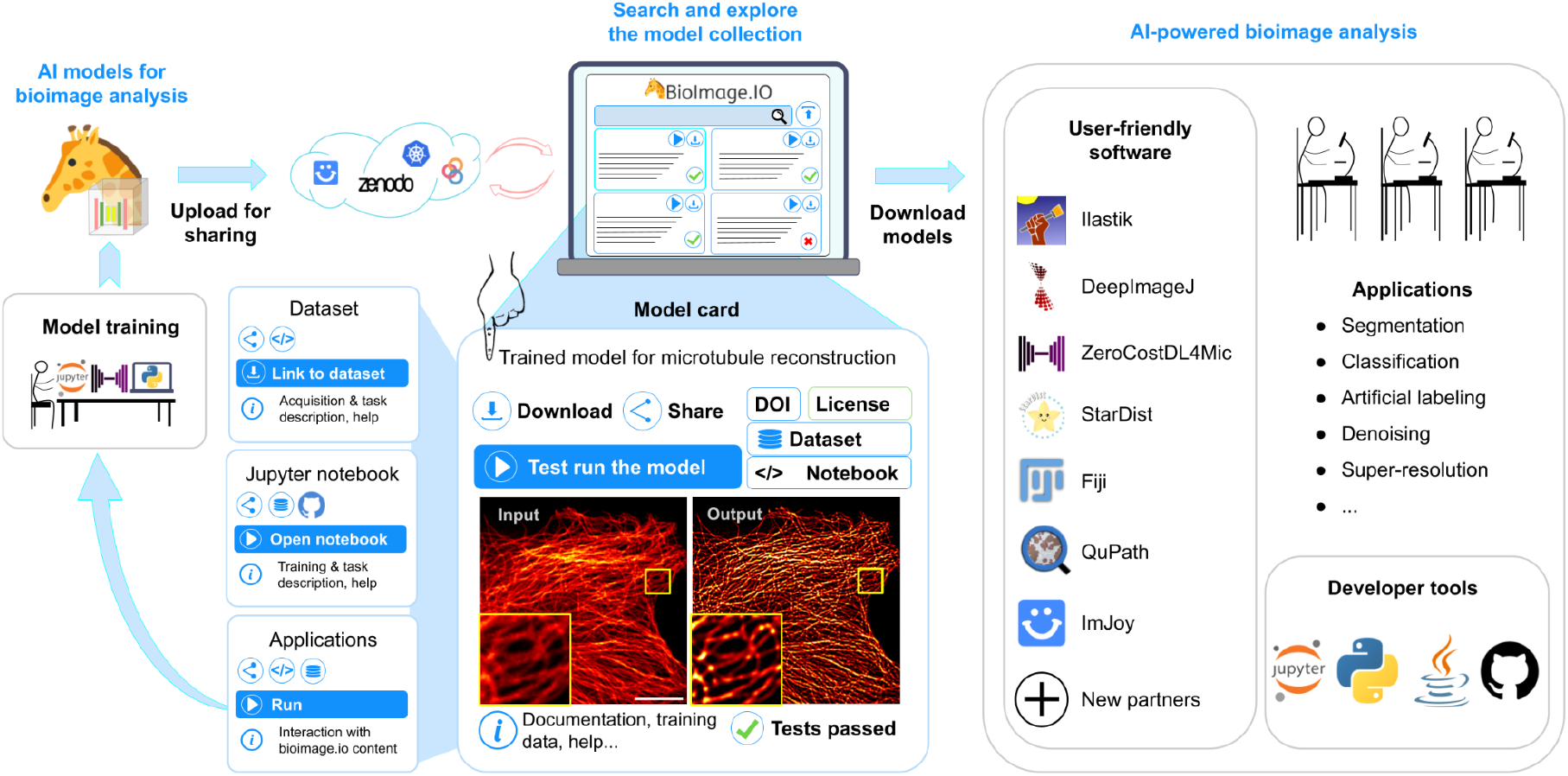
Life-cycle of deep learning models in the BioImage Model Zoo. A trained model following the format specifications in the Zoo is uploaded to bioimage.io through the web interface and its content is hosted in Zenodo. Any user can search and download any model in the BioImage Model Zoo. They can apply the model to analyze new data in the user-friendly software supported by community partners. Furthermore, trained models can be directly integrated in custom-made bioimage analysis workflows by using our Python or Java libraries.. Each model in the BioImage Model Zoo is displayed through an interactive model card that provides core information (e.g., description, license, authors and publication information, source code, and links to the training datasets and notebooks to (re-)train the models) and enables interaction with its content (i.e., processing an image with the trained model in the browser). Finally, any user can (re-)train or fine-tune the model, and contribute it back to the BioImage Model Zoo.

To life scientists and method developers alike, we offer the bioimage.io website, built to provide an interactive, user-friendly experience with all the content in the BioImage Model Zoo. Our growing collection already contains models for a variety of popular microscopy image analysis tasks. Importantly, models of the Zoo are not stored in isolation: to enhance reproducibility, we store references to the associated datasets and model creating notebooks. Additionally, pre-trained networks combined with notebooks that trained these networks are providing an ideal set of tutorial-like basis for non-expert users to see and learn how they can train similar models on their own data. In this way every user who wishes to train new models can learn to do so and contribute their trained models to the Zoo. Furthermore, inspired by previous approaches^12,28,29^ for browser-based interaction with DL models, the Zoo is enriched with the ImJoy application framework called the *BioEngine* which enables users to test models on their own data directly in the browser.

All models in the Bioimage Model Zoo can be searched and downloaded for future use in a growing number of desktop- and cloud-based tools, and referenced by their unique identifier when mentioned in scientific works, hence making this new infrastructure adhere to the FAIR^35^ principles. We envision that the BioImage Model Zoo will become a big step towards democratizing access to the latest AI developments in the life sciences.

## Results

### Standardized DL models

Interoperability between desktop tools has always been a significant concern for the bioimage analysis community. Unavoidable incompatibilities introduced by programming language and platform heterogeneities have challenged the community and limited global dissemination efforts and direct tool interoperability. While DL comes with specific demands and requirements, we believe it offers the unique opportunity to bring computational tools closer together: many pre-trained networks contain all the necessary information to apply them to new data. A well-defined, open specification of the pre-trained network metadata will allow performing this step programmatically, *i.e.,* without the need to read and modify the underlying network source code. In the BioImage Model Zoo, we have defined such a specification following the requirements of initial community partners (see Online Methods). While the format is designed with flexibility in mind, we expect it to evolve further with the development of DL tools, frameworks and with more community partners joining our efforts. Importantly, our specification is a syntactic wrapper of the existing formats for neural network weights, which enables direct programmatic network inference on new data. The weights themselves are stored as defined by the corresponding DL frameworks, e.g., PyTorch or Tensorflow. Starting from the current version (v0.4), we commit to keeping our infrastructure and libraries backward-compatible to future format amendments, i.e., all contributed models will remain usable.

Every model in the Zoo has a DOI (see Model Submission) and is additionally assigned a nickname that is easier to remember, composed of an adjective and an animal name, e.g. *chatty-frog* or *passionate-t-rex*. The BioImage Model Zoo format stores all of the network metadata as required or optional fields. Required fields contain the technical specifications of the model that are necessary for correct deployment (e.g., input and output shape, pre- and post-processing routines). Additionally, there are required fields that are not strictly necessary for running the model but they ensure full reproducibility and proper credit attribution within the Zoo and in derived work (*e.g.*, “Author”, “Contributor”, or “License”). In contrast, a plethora of additional useful information can be added to integrate specific metadata needed by diverse DL-based workflows, *e.g.*, StarDist^25,26^ (see Figure 2). Thus, the community partners support all required fields of the BioImage Model Zoo format in their corresponding tools and introduce their optional fields for more complex workflows or in anticipation of future needs. We work towards the vision of making all trained neural networks in the Zoo executable in all the open source tools represented by the community partners, barring the intrinsic incompatibilities between deep learning frameworks.

**Figure 2.**
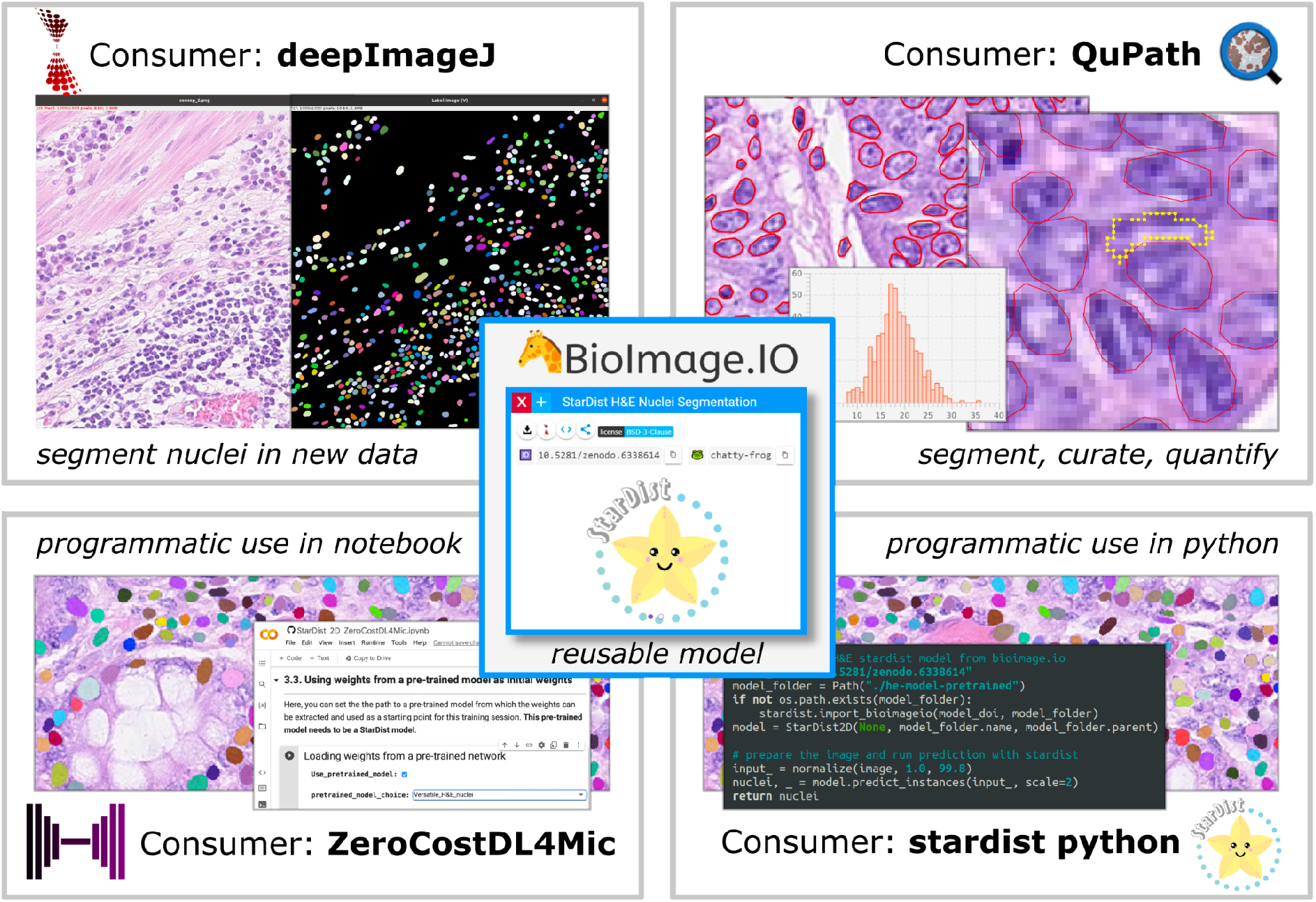
Segmentation of nuclei in H&E stained tissue sections. (Center) The pretrained StarDist model for nucleus segmentation in H&E stained images is available in the BioImage Model Zoo. We run it in four different consumer softwares that support the model format and StarDist specific post-processing: (top left) deepImageJ, which allows the integration with the StarDist Fiji plugin and the Fiji ecosystem; (top right) QuPath, which provides convenient curation and quantification; (bottom left) ZeroCostDL4Mic, which also offers fine-tuning of the model on new data; (bottom right) StarDist Python library, which enables integration within Python analysis pipelines.

### A user-friendly resource for the whole community

The BioImage Model Zoo provides a unified and accessible DL models repository to promote their use across different bioimage analysis tasks. On the BioImage.IO website, every model is displayed as an auto-generated interactive “model card” (see Figure 1). It shows the example input-output data pair and lists the most critical information about the model, such as the author(s) and instructions on how it can be run and validated. Most model cards allow users to open the in-browser “Test Run” application and test the model on example images as well as on their own data. Furthermore, each card provides a channel for communication with users and encourages them to leave (moderated) comments about the model. In addition to the pre-trained model weights and metadata, the model card can keep a link to the training data and notebooks. To make these persistent, we support very basic cards to describe notebooks and datasets hosted elsewhere, they can be found under “Applications” and “Datasets” tabs of the website. We strongly encourage our contributors to provide the full chain of training data, model weights and the training code.

As an illustration of the analyses we aim to enable, here we describe four microscopy use cases that cover the range from directly applying a pre-trained model, fine-tuning a model without using any code, incorporating several models into custom applications, and publishing a recently developed method for domain adaptation.

#### Use case 1: Applying pretrained models across tools (Fig. 2)

Here, we consider an example problem of nucleus segmentation in H&E stained tissue sections. We choose the pre-trained model for the H&E modality from the popular instance segmentation method StarDist^16,17^. This model was trained on two H&E datasets^38,39^ using the StarDist Python library and is versatile enough to be applied to similar yet slightly different H&E data. This model is now available in the BioImage Model Zoo (id: 10.5281/zenodo.6338614 or *chatty-frog)* and can be deployed in multiple supporting tools. Specifically, we use this model to segment H&E images from the recently released Lizard dataset^40^ in deepImageJ, QuPath, ZeroCostDL4Mic, and the StarDist Python library (see Fig. 2). Through the integration with all of these tools, researchers can now choose to apply the model with the tool they are most familiar with or that is most suitable for their further analysis needs, for example by enabling integration with other Fiji plugins (deepImageJ), convenient curation and quantification (QuPath), fine-tuning of the model on new data (ZeroCostDL4Mic) or programmatic use in Python (StarDist Python library).

#### Use Case 2: Easy fine-tuning (Fig. 3)

In this case study, we demonstrate how a model from the BioImage Model Zoo can be executed in ilastik, fine-tuned in ZeroCostDL4Mic and executed again in deepImageJ. We address the problem of segmenting tissues into cells based on cellular membrane staining. For this, the DL models corresponding to the state-of-the-art solution^41^ are available through the BioImage Model Zoo (id 10.5281/zenodo.6334583, or *passionate-t-rex*) and the training procedure has been reimplemented using the ZeroCostDL4Mic framework (id 10.5281/zenodo.5749843 or *humorous-owl)* (see Figure 3). As the first step, we download the model and run it in the ilastik Neural Network Classification workflow on a publicly available dataset that was not part of the training set^41,43^. We use the boundary predictions in the ilastik Multicut workflow to correct a few of the wrong segmentation edges and build a new annotated dataset to fine-tune the model in the corresponding ZeroCostDL4Mic notebook (see Figure 3a). The fine-tuned model – well adapted to the new dataset – is exported from the notebook and uploaded to the BioImage Model Zoo (id 10.5281/zenodo.6348728 or *non-judgemental-eagle*) (see Figure 3b). Finally, the model can be used in deepImageJ as part of a larger analysis pipeline, including also morphological measurements (see Figure 3c). The original training data, also displayed in the BioImage Model Zoo, is linked to the trained models through the metadata listed in the model description file. Thus, anyone can try to improve the performance of the segmentation by incorporating the most recent computer vision ideas.

**Figure 3.**
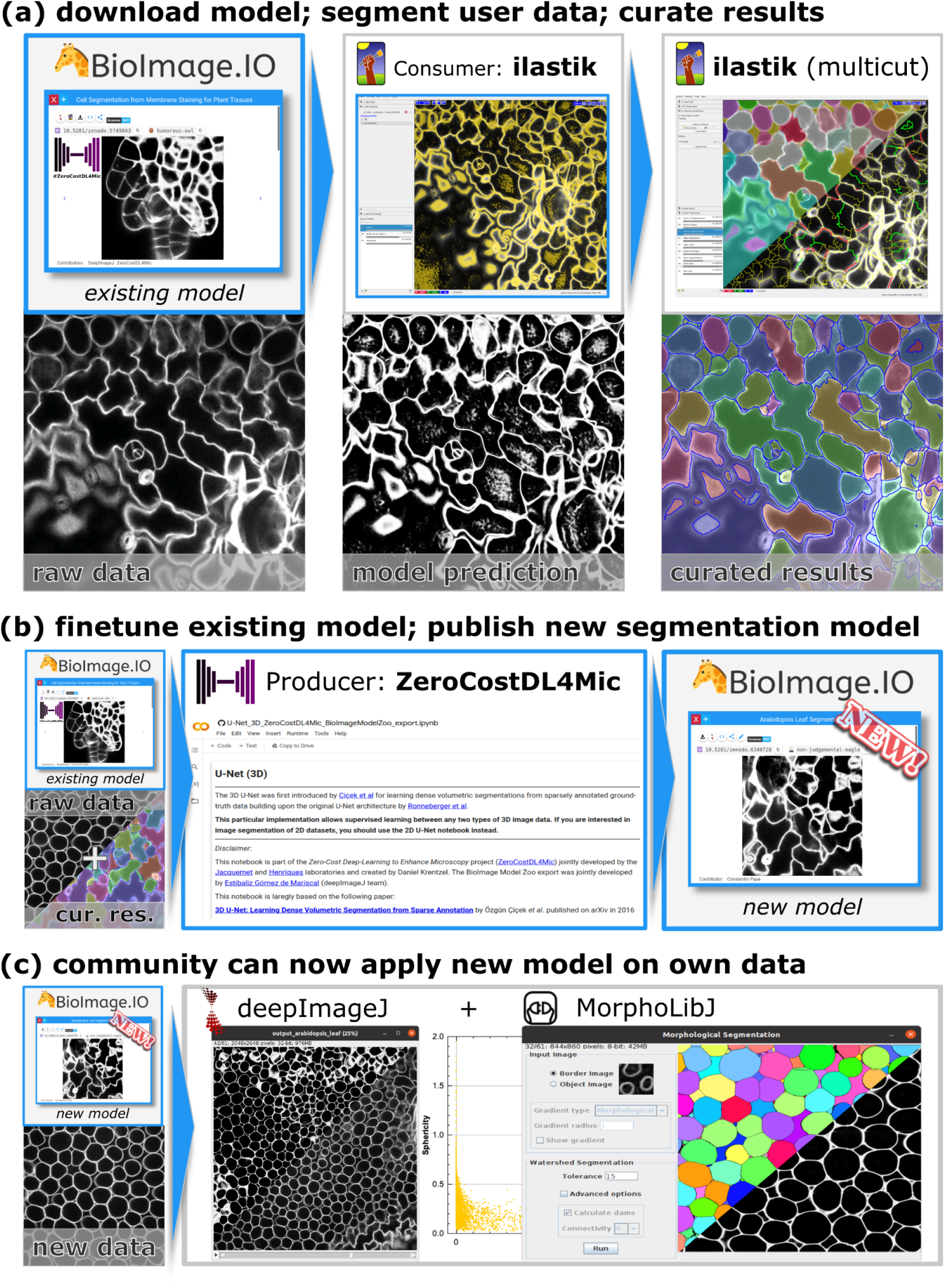
Boundary-based segmentation of cells in light microscopy; (a) The 3D U-Net for cell segmentation in confocal stacks of *Arabidopsis thaliana* ovules is available in the BioImage Model Zoo. We apply it to an independent dataset featuring arabidopsis leaves^41^. The predictions are very noisy, complicating direct post-processing. We generate a curated instance segmentation using the predictions as input to the interactive ilastik multicut workflow for boundary based segmentation; (b) We fine-tune the model on the curated segmentation from (a) in ZeroCostDL4Mic, obtaining a model that performs better on the new data. We upload the new model to the Zoo; (c) The new model is now available for everyone to use. Here, we apply it to the leaf data using deepImageJ and then obtain an instance segmentation through simple watershed post-processing with MorphoLibJ^42^, which enables us to extract per cell measurements such as sphericity distribution.

#### Use Case 3: Building multi-model pipelines (Fig. 4)

In this example, we take the winning DL solution from the Human Protein Atlas (HPA) single cell classification competition hosted on Kaggle (https://www.kaggle.com/c/hpa-single-cell-image-classification/) and make it available in a BioEngine web application and a standalone application implemented with napari^32^. The challenge addressed both cell segmentation and classification of cells according to the protein localization pattern. The proposed workflow starts by running a UNet-based model for cell and nucleus segmentation to obtain the cell masks. Then, each masked single cell image is fed into another DL model that predicts one or several protein localization labels. We have uploaded the three models for cell boundary prediction (id 10.5281/zenodo.6200635 or *loyal-parrot),* nucleus prediction (id 10.5281/zenodo.6200999 or *conscientious-seashell)* and protein classification (id 10.5281/zenodo.5910854 or *straightforward-crocodile)* to the BioImage Model Zoo.

We use the bioimageio.core Python library (see Online Methods) to implement a desktop application that downloads the three models, performs cell segmentation and classification as well as visualization of the results using napari (see Figure 4a). Similarly, we use the BioEngine to implement a web-based application that performs segmentation and classification online using the models in the Zoo, and visualizes the results (see Figure 4b). The BioEngine app can be started straight from the app icon on the model card and runs directly in the browser. In either case, only a minimal amount of simple Python code is needed to integrate a pipeline of multiple networks with a user-friendly GUI.

**Figure 4.**
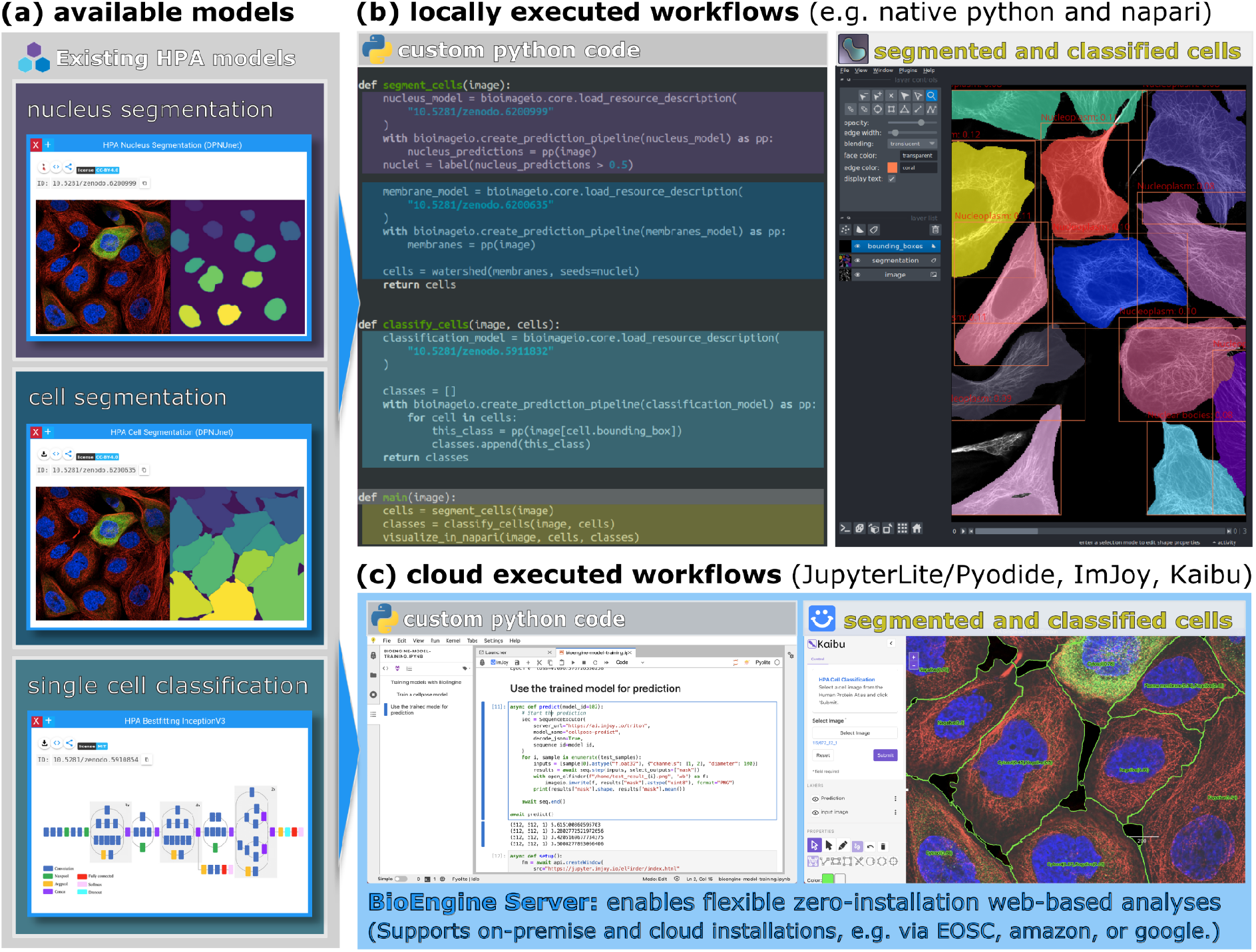
Building a single cell classification analysis workflow with three models in the BioImage Model Zoo, executed locally using our Python library or through the BioEngine. (a) Models for the winning solution of the Human Protein Atlas single cell classification competition are hosted in the BioImage Model Zoo. Images from the Human Protein Atlas were first segmented with the nucleus segmentation and cell segmentation models, and for each segmented cell, an InceptionV3-based model predicts the protein localization, based on the green channel. (b) The workflow can be reproduced using our “bioimageio.core” python library locally, using napari to visualize the results. (c) With the BioEngine it can also be replicated directly in the browser, using Kaibu (an ImJoy plugin) for visualization. The BioEngine application executes models on remote servers (on-premise or in the public cloud) and the workflow can be composed in a JupyterLite notebook or in an ImJoy plugin.

#### Use Case 4: Dissemination of new DL approaches (Fig. 5)

Here, we show how the pre-trained models of the Zoo can be exploited for domain adaptation through the approach proposed by Matskevych et al^44^. We use their trained model for mitochondria segmentation in electron microscopy (EM) already available in the BioImage Model Zoo (model id: 10.5281/zenodo.6406756 or *hiding-blowfish)* and the images from the MitoEM challenge^45^. This model is trained to improve 2D mitochondria predictions from a “shallow” classifier *(e.g.,* Random Forest). This approach provides more robust results for mitochondria segmentation in EM modalities when applied to new images since the model only sees the changes in the shape distribution, rather than the intensity distribution. For a new EM dataset, “shallow” pixel classification can be performed fast and interactively using established tools such as ilastik, LabKit^46^ or the Trainable Weka toolkit^47^. The “enhancer” model can be applied using the ilastik neural network workflow or deepImageJ to significantly improve the segmentation results without further data annotation or training.

**Figure 5.**
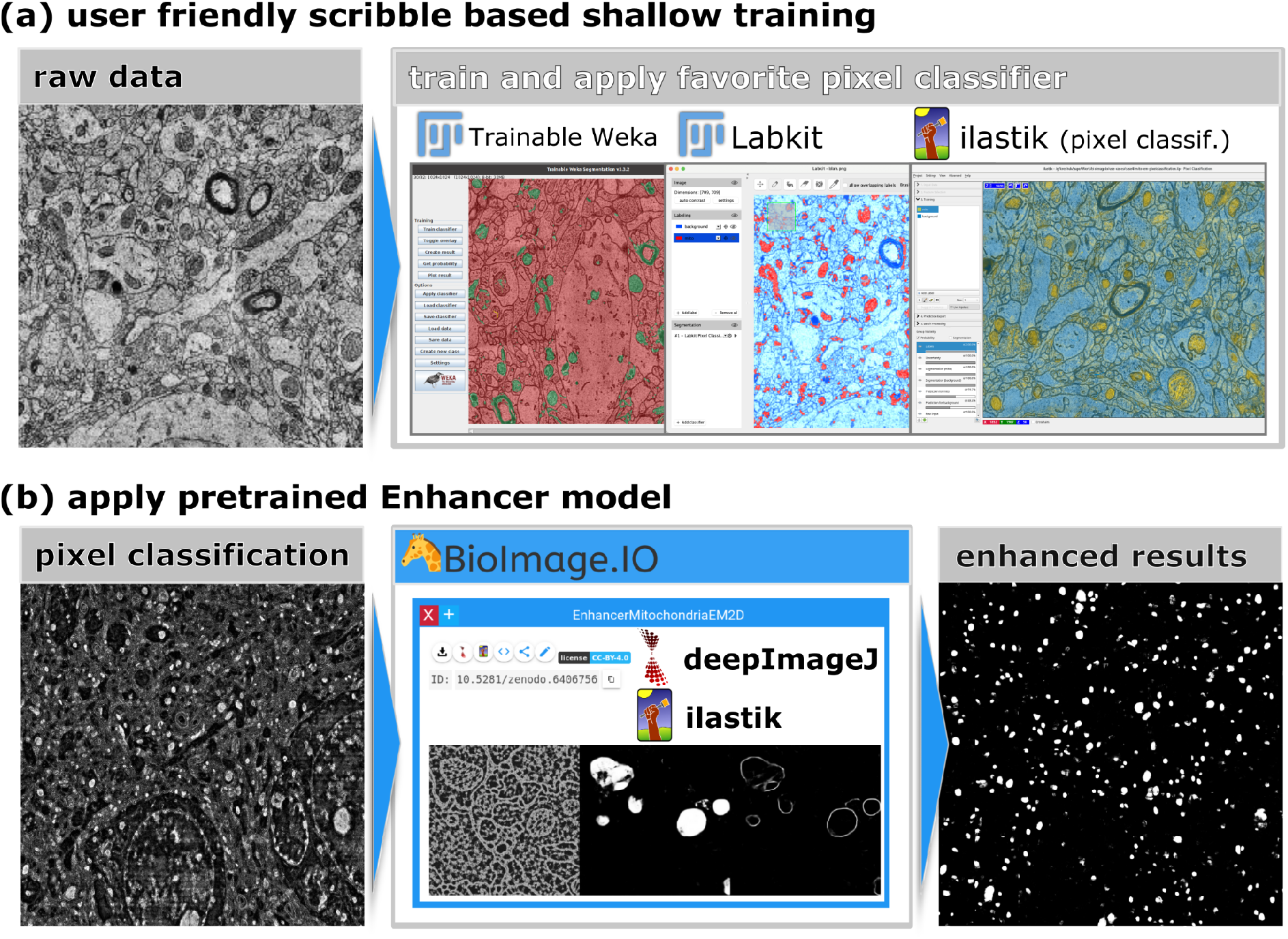
Domain adaptation for EM mitochondria segmentation with pre-trained models from the Zoo; (a) We show several options for interactive training of a shallow classifier (i.e., Random Forest) that segments mitochondria in new EM data: Fiji plugins “Trainable Weka segmentation” and “Labkit’’ and the pixel classification workflow in ilastik; (b) The domain adaptation model trained with the method from Matskevych et al^44^. is available in the Zoo. It can be used to significantly improve the mitochondria predictions from the Random Forest classifier, using either deepImageJ for the results obtained with Weka/Labkit or the ilastik neural network workflow for the ilastik pixel classification result.

### A growing network collection

The BioImage Model Zoo contains trained DL models for popular microscopy image analysis tasks, from nucleus segmentation in light microscopy over modality agnostic image restoration methods, to membrane segmentation in electron microscopy images. It is aimed for anyone to submit new ones independent of the bioimage analysis application. We benefit from the version traceability and DOI assignment of Zenodo (https://zenodo.org/) to store models. Contributors are guided through every step of the submission process by a detailed tutorial (https://bioimage.io/docs/#/contribute_models/). The intellectual property of each model is fully preserved with the information of the original authors, who also decide on the license of model re-use. Any new contribution is expected to (*i*) conform to the defined metadata format, (*ii*) provide a test image for automatic testing and *(iii)* contain sufficient documentation on how the results can be validated. Each new model submission triggers automatic quality assurance which is finished by the manual acceptance or rejection of the model by a human curator.

The BioImage Model Zoo integrates programmatic tools for Python and Java that allow developers to import, execute, and export specification-conforming models. Furthermore, we build on the success of the ZeroCostDL4Mic project to simplify the training, testing and use of DL models and equip some of the most used notebooks (i.e., 2D and 3D U-Nets, 2D U-Net for semantic segmentation and 2D StarDist) with a final step to export the trained model to the BioImage Model Zoo model file format, ready for direct upload to the Zoo.

Because DL models are critically dependent on the training schedules and datasets, and intrinsically on the scripts used to train them, the BioImage Model Zoo also reserves space to gather information about datasets and Python notebooks. Our metadata format includes fields for training data and general links, e.g. for training notebooks, while the Zoo itself can host links to datasets and notebooks in an interconnected manner. Different models can be linked to the same dataset or notebook, supporting the tracking of models that improve the performance of a specific task or allowing researchers to upload new models that share their architecture but have been trained differently.

The performance of a DL model strongly depends on its training data. Since networks only learn from the images in the training set, they are specialized to perform well on similar images, but not every network will perform equally well on user’s data. For a quick and easy qualitative validation of the models in the Zoo, the BioEngine enables model testing directly from the bioimage.io website. For most models, users can upload an example image or even a small dataset and process it automatically in one click in the browser or via our cloud server infrastructure (de.NBI cloud). We rely on model developers to provide sufficient documentation on how their model can be validated. Additionally, every model is required to contain a test input image and the corresponding expected output. These are used within our continuous integration pipeline to test the model by automatically comparing the obtained prediction with the provided test output. The documentation for all existing models in the Model Zoo contains a section on “Recommended Validation”, describing the appropriate metrics and often also contains links to scripts or example notebooks.

With these simple submission tools and multiple training examples, we hope to enable image analysts outside the narrow circle of professional DL method developers to contribute their models to the community (see Use Cases 1 and 2 and Figure 1).

### Web application for testing models

To further facilitate the adoption and evaluation of DL methods among non-expert users in the life sciences, we provide point-and-click “Test Run” buttons for quick execution and evaluation of the models in the BioImage Model Zoo via a ready-to-use web application. More specifically, we built an application framework, named BioEngine, on top of the ImJoy plugin framework and recently developed cloud computing infrastructure for AI model serving. The BioEngine backend is a multi-model and multi-user model serving framework built on top of the Nvidia Triton Inference Server (https://developer.nvidia.com/nvidia-triton-inference-server). It allows mounting the entire repository of models of the BioImage Model Zoo on the server, and dynamically scheduling the model inference tasks into a limited number of GPUs based on users’ requests. Like DeepCell Kiosk^28^, we use a Kubernetes cluster to manage a set of container-based server components and perform auto-scaling of the computational instances in response to the varying number of users. The BioEngine allows the user to load their image files via a user-friendly ImJoy web application, to be processed either in the browser or on our remote cloud servers. The result is then displayed directly in the browser using image viewers such as Kaibu^49^, itk-vtk-viewer^50^, or vizarr^51^ (see also Use Case 3 and Figure 4). We provide the “Test Run” BioEngine application to enable basic interaction with most models in the browser. Model contributors can create more tailored applications for each model to improve the user experience further or combine models in a workflow (as demonstrated in Use Case 3 and Figure 4). More examples, such as the Mutex Watershed^52^ plugin for affinity-based instance segmentation (id: 10.5281/zenodo.6079314 or *wild-whale*), can be found in the Zoo.

While these applications and computing infrastructure are meant for demonstration and evaluation purposes on small amounts of data, we envision a future where such infrastructure can be distributed for on-premise deployment and maintained by experts in users’ research institutions and universities.

## Discussion and Outlook

We introduced the BioImage Model Zoo: a community-driven infrastructure which is designed to *(i)* enable easy and direct use of cutting-edge AI for life science applications in a growing number of user-friendly open-source software tools, (*ii*) promote the rapid and effective dissemination of DL developments for bioimage analysis, and *(iii)* enrich and support advanced bioimage analysis workflow development by enabling the interoperability of DL models.

The BioImage Model Zoo aims to become the key repository where DL models can be deposited and shared according to FAIR principles:

- (Findable) All our models are freely available and displayed through a convenient search interface of the Zoo, uniquely identified with a DOI and a memorable nickname.
- (Accessible) For model users, we provide a rapidly growing collection of pre-trained models that can be used from within multiple popular “point-and-click” analysis tools created by members of the BioImage Model Zoo community. We envision that integration with “point-and-click” tools will greatly simplify collaboration between biologists and method developers or analysis facility staff. By simply filling out the fields of our model metadata format, the developers already make their model accessible to non-computational collaborators.
- (Interoperable) We provide a flexible open network format specification and libraries for developers that streamline the creation and consumption of models in all types of user-friendly and programming tools.
- (Reproducible) The training history of each model and the provenance of its training data can seamlessly be traced back through the model description format we proposed. Furthermore, we encourage model developers to link the models to available training code and openly accessible data in the form of runnable scripts or notebooks. Integration with user-friendly training tools such as ZeroCostDL4Mic enables the users of such tools to become model contributors and in this way, paves the way for open and FAIR science in the age of DL-based image data analysis.

The BioImage Model Zoo is tailored for non-expert users willing to exploit DL in their routine image analysis tasks. We designed our web-based model repository in a user-friendly spirit, with models displayed as documented graphical cards with interactive buttons which let any user dive into the model content. The test images and outputs provide a quick view on the applicability of each model to a new user task, while the test run functionality allows to evaluate it more precisely. We envision that the BioEngine – the technology behind the test runs – will in the future be expanded to provide more web-based tools associated with the Zoo and its models, such as the example segmentation plugin we describe above.

We have designed the model metadata format with a focus on interoperability and reproducibility. Still, due to the stochastic nature of network training procedures, no full reproducibility can be achieved even when the same code and the same training data is used. Similarly, no theoretical guarantees can be given on the performance of any network on previously unseen data. We hope that future research in Bayesian deep learning, uncertainty estimation and out-of-distribution sample handling will allow the community to address these challenges in a more general and principled manner. For the moment, we rely on model contributors to recommend the validation workflows most appropriate for the image analysis task.

AI-based methods have the critical potential to boost life sciences research through automation of all image-related tasks, from smart microscopy to novelty detection and quantitative image analysis at scale. The BioImage Model Zoo provides prompt access to novel models that are packaged for easy use in popular image analysis tools, scripts or custom pipelines. Through the integration of ZeroCostDL4Mic and, in general, the development of easy-to-use libraries for running the models in scripts and notebooks, we hope to turn many life scientists into the Zoo contributors, sharing (re-)trained models and training datasets for the benefit of the whole community.

Finally, the benefits to the method development community will not be limited to the improvement of the deployment process. A collection of cross-compatible models will allow for multi-component modular pipelines as shown in Use Case 3, where each component can be optimized separately or jointly with others. Availability of multiple models trained for the same task will enable building model ensembles, improving robustness and generalization of the final prediction. Furthermore, such model collections will give a new boost to research on automatic model selection, ultimately making the Zoo even more accessible to the whole microscopy community. We expect this type of interdisciplinary exchange to become a critical step towards accelerating and advancing life sciences research.

## Acknowledgments

We thank Jean-Christophe Olivo-Marín, Stéphane Dallongeville, and Jean-Yves Tivenez for their valuable input and feedback about building synergies between bioimage analysis software. We also thank Steffen Wolf for helping with the integration of the Mutex Watershed Segmentation plugin. We would also like to thank the EMBL-EBI training programme and ZIDAS Image-Analysis for Biologists course organizers for letting us show our work and receive constructive feedback from life scientists during the development of the BioImage Model Zoo. We also thank Romain Laine for supporting the update of ZeroCostDL4Mic and providing important feedback to the BioImage Model Zoo specifications. This work was supported by the BMBF-funded de.NBI Cloud within the German Network for Bioinformatics Infrastructure (de.NBI) (031A532B, 031A533A, 031A533B, 031A534A, 031A535A, 031A537A, 031A537B, 031A537C, 031A537D, 031A538A) and the European Commission through the Horizon Europe program (AI4LIFE project, grant agreement 101057970-AI4LIFE). A.K., M.N., C.P. and D.K gratefully acknowledge the support of the Deutsche Forschungsgemeinschaft (DFG) under grant HA-4364/11-1. W.O. and E.L. would like to acknowledge the support of the Knut and Alice Wallenberg Foundation KAW2018.0172 and 2021.0346 to EL) and the Swedish Research Council (2018-06461_3 to E.L.). This work was also also supported by Ministerio de Ciencia, Innovación y Universidades, Agencia Estatal de Investigación, under Grant PID2019-109820RB-I00, MCIN / AEI / 10.13039/501100011033/, co-financed by European Regional Development Fund (ERDF), “A way of making Europe” (EGM, grant to A.M.B.); BBVA Foundation under a 2017 Leonardo Grant for Researchers and Cultural Creators (A.M.B.). E.G.M, C.G.L.H., L.M.S., C.T.G. and A.M.B. acknowledge NVIDIA Corporation for the donation of the Titan X (Pascal) GPU (to AMB). E.G.M. and R.H. are supported by the Gulbenkian Foundation, European Research Council (ERC) under the European Union’s Horizon 2020 research and innovation programme (grant agreement no. 101001332) and the European Molecular Biology Organization (EMBO-2020-IG-4734). R.H. is further supported by a Chan Zuckerberg Initiative Visual Proteomics Grant (vpi-0000000044). G.J. is supported by the Finnish Cancer Organization (grants to G.J.), Sigrid Juselius Foundation (grants to G.J.), Academy of Finland research fellowships (338537 to G.J.), the Åbo Akademi University Research Foundation (CoE CellMech; to G.J.), the Drug Discovery and Diagnostics strategic funding to Åbo Akademi University (to G.J.). M.W. is supported by the EPFL School of Life Sciences and a generous foundation represented by CARIGEST SA. Deb.S. was funded by Helmholtz Imaging, a platform of the Helmholtz Incubator on Information and Data Science.

## Online Methods

### Community Partners

While the BioImage Model Zoo is open to contributions from anyone, some groups, e.g., particular GitHub organizations, may contribute extensively or seek a closer integration of their software and the BioImage Model Zoo. Often these groups can also be granted a credit of trust as they contribute to maintenance and development efforts. Our current community partners are ilastik, deepImageJ, QuPath, StarDist, ImJoy, ZeroCostDL4Mic, CSBDeep and HPA. Community partners are listed on the bioimage.io website; unlike other contributors they are allowed to contribute resources without manual approval (see Contributing to the BioImage Model Zoo section in the Online Methods). Furthermore, their continuous integration scripts can be included in (model) resource testing (see Model testing infrastructure in the Online Methods). Any group can become a community partner simply by creating a GitHub issue stating their intentions and supplying some basic information on their project. Upon approval, the subsequent steps include creating their own resource collection. In more detail, we set up a GitHub Actions workflow to validate their collection, for which we provide templates, and optionally, another GitHub Actions workflow to test the functioning of their software with the BioImage.IO resources. This is described in more detail in the following sections (see Contributing to the BioImage Model Zoo and Model testing infrastructure in the Online Methods). A step-by-step guide is provided as part of our updated documentation at https://bioimage.io/docs/#/community_partners/README.md.

### The bioimage.io website

The bioimage.io website is the central entry point for users of the BioImage Model Zoo. It enables users to browse the bioimage.io resources, provides a user-friendly upload form to contribute such resources and hosts our documentation (https://bioimage.io/docs). The website’s source code and our documentation are available at https://github.com/bioimage-io/bioimage.io. The website’s content –the BioImage.IO resources– are fetched dynamically from the bioimage.io collection (details in Resource hosting and serving in the Online Methods). Each resource has a unique ID. For resources hosted via Zenodo, the DOI of the entry is used as the ID. For resources contributed directly via a community partner, the ID is specified in the partner repository and prefixed with the partner name in order to guarantee uniqueness. Models have an additional associated animal nickname in order to provide a more memorable identifier. Contributed models need to provide example test input and output which are used by the Zoo’s infrastructure to perform automatic quality assurance, reducing the chances of failure when deploying them in the consumer software. We strongly encourage resource contributors to provide extensive tags that enable efficient model search and discovery in the Zoo. To this end, we provide a library of tags following the EDAM ontology^48^ and use those to enable search by keywords or free text.

Each resource is represented by a card displaying compact information such as name, a short description, cover images and links to other bioimage.io resources *(e.g.,* link to the notebook to train a model and vice versa). Links to applications provide user-friendly interaction with the resource via BioEngine Apps (see “BioEngine”). For example, each model is linked to the bioimage.io *Packager* application, which allows the download of ready-to-use models. The card representing a bioimage.io resource expands on click for more details–full description, contributors and citations–and convenient buttons to copy the resource ID or animal nickname. These identifiers can be used to refer to the model in consumer software provided by the community partners, custom scripts/notebooks utilizing our developer tools (see Developer tools in the Online Materials) or for documentation purposes. Extended details for models also include a description of model training, as well as test summaries reporting which software (versions) can run inference with the given model (see Model testing infrastructure in Online Methods).

### Resource Types & Formats

BioImage Model Zoo can store and display multiple kinds of Resources, described through Resources Description Files (RDF) which are stored as a YAML file. Potential Resources include models, datasets, notebooks and applications, but we envision other types of Resources arising with further development of the Zoo. The “general RDF” contains the most essential information we require to be able to build a card for a Resource and show it to our users. For models and collections, additional metadata information is expected to enable the Zoo testing functionality.

### General RDF

A (general) RDF is a YAML file that adheres to our RDF specification. The required and optional fields in an RDF, such as name, description, authors, or citations are described in detail in our documentation (https://bioimage.io/docs/#/bioimageio_spec).

To empower the bioimage.io consumer software, we enable arbitrary fields (gathered within the *config* field) to specify additional metadata information, which is not (yet) incorporated into our RDF specification but that enables further exploitation of the content of the Zoo.

For some resource types–models and collections, at the time of writing–this general RDF is extended to provide the minimal technical information required to ensure the cross-compatibility and deployment in the consumer software.

#### Model RDF

The Model RDF (https://bioimage.io/docs/#/bioimageio_model_spec) extends the general RDF specification to describe trained neural networks. It has a ‘weights’ field that contains the location of one or several network weight files *(i.e.,* weights of a model instance stored in different formats). Currently, we support the following weight formats: Keras HDF5, ONNX, PyTorch state dictionary, Tensorflow Javascript, Tensorflow Saved Model Bundle and Torchscript (see https://bioimage.io/docs/#/bioimageio_weights_spec). The input and output tensors of the model are described in additional fields. These ‘inputs’ and ‘outputs’ fields hold a description of the data type, a shape description and the axis types and order, which is restricted to up to three spatial axes (‘xyz’), a channel axis (‘c’) and a batch axis (‘b’). Each input/output tensor has its own ‘description’ text and may specify transformations–’preprocessing’ for ‘inputs’, ‘postprocessing’ for ‘outputs’–from a defined set (https://bioimage.io/docs/#/bioimageio_preprocessing_spec, https://bioimage.io/docs/#/bioimageio_postprocessing_spec). Fields for test inputs and outputs provide data for testing that the model runs as expected. The optional sample inputs and outputs can be used to illustrate model use hands-on. Model RDFs additionally contain documentation on how the model was trained. Like any RDF, a model RDF may be linked to other bioimage.io resources such as notebooks and datasets, which enable experienced users to reproduce or refine the model.

#### Collection RDF

Collection RDFs conveniently describe a group of resources and are primarily used by bioimage.io community partners to contribute resources via GitHub to the BioImage Model Zoo. The collection RDF extends the general RDF by a ‘collection’ field holding a list. Each list entry represents an independent RDF, which is based on the collection RDF itself without the ‘collection’ field, updated first by content loaded from the entry’s ‘‘rdf_source’ field if present, and second, by any other field specified in the entry, except for the ‘id’ field. The collection needs to have an ‘id’ field (*e.g.,* id: ‘ilastik’) and the same for each entry *(e.g.,* id: ‘torch-em-3d-unet-notebook’), which expand to a unique identifier in the BioImage Model Zoo for this specific resource *(e.g.,* id: ‘ilastik/torch-em-3d-unet-notebook’). Therefore, each collection entry is required to have a unique identifier. The prepending of the collection id guarantees that nested collections have unique resource identifiers as well.

### Resource hosting and serving

All repositories that comprise the BioImage Model Zoo are hosted on GitHub. Our documentation (https://bioimage.io/docs/) utilizes the ImJoy Docs tool (https://imjoy-team.github.io/imjoy-docs/) and is hosted with the bioimage.io website source at https://github.com/bioimage-io/bioimage.io.

The BioImage Model Zoo content displayed by the website is managed by the bioimage.io collection repository (https://bioimage.io/docs/#/bioimageio_collection_repo), which holds a curated list of resources. The bioimage.io resources defined by the RDF are either hosted on Zenodo or in dedicated GitHub repositories in the respective partner GitHub organization (see “Contributing to the Model Zoo”).

### Contributing to the BioImage Model Zoo

The default way to contribute a model is to upload the model RDF file *(i.e.,* ‘rdf.yaml’) to a Zenodo record linked to the bioimage.io community in Zenodo.

Alternatively, the bioimage.io website is equipped with an interactive guide to upload the model metadata and all the required files. This process validates the metadata directly in the browser using our bioimageio.spec library (see “Developer tools’’). Then, it publishes the new resource on Zenodo under the contributors Zenodo username. This process also triggers an update of the bioimage.io collection via a netlify server to avoid any delays in the process of finalizing the contribution as described below.

Note that our core library can be used to create a bioimage.io model programmatically with the ‘build_model’ function (see Developer tools in the Online Methods).

Updating the bioimage.io collection is facilitated by a GitHub Actions workflow in the collection repository. This workflow queries Zenodo for records with the ‘bioimage.io’ keyword and generates a pull request (PR) in the BioImage.IO collection repository for each new resource. Such a contribution PR adds a resource entry in the form of a ‘resource.yaml’ file based on the ‘rdf.yaml’ file in the Zenodo record and serves as a platform for bioimage.io maintainers and contributors to discuss the inclusion of the new resource. Relevant aspects and the conclusion of this discussion are described in the following paragraph.

Upon creation of a contribution PR, a generated animal nickname, e.g. ‘creative-panda’, is added to the ‘resource.yaml’, which serves as an alternative, more memorable resource identifier. The generated animal nickname may be altered in the first contribution PR for a given resource. Additionally, any field of the original ‘rdf.yaml’ may be updated, which is intended for small changes like typos or formatting. Substantial changes should be reflected in the Zenodo record as a new version.

A Zenodo record version has a concept DOI and a version DOI (https://help.zenodo.org). All versions of a Zenodo record share the same concept DOI. In the bioimage.io collection repository each ‘resource.yaml’ corresponds to the concept DOI of a Zenodo record and holds a list of the existing versions. A resource as a whole or a particular version of it may be blocked by setting a ‘status’ field in the ‘resource.yaml’ to ‘blocked’.

For each accepted version, an updated ‘rdf.yaml’ is generated and validated by a GitHub Actions workflow running in the contribution PR to ensure that the resource is a valid RDF and that test outputs can be reproduced from test inputs. In addition to this technical check, a bioimage.io maintainer checks that the resource has an intuitive name and a suitable description, as well as complete documentation. Once these requirements have been satisfied, a bioimage.io maintainer will merge the PR and accept the new resource (version).

Yet another option to contribute is through a community partner. This implies adding or updating a resource to the community partner’s registered collection RDF. For the time being, model resources may only be contributed via Zenodo to ensure their persistence.

To update a contributed resource a user can create an updated version of the respective Zenodo record (either through our interactive upload guide or manually on zenodo.org). This will trigger a PR analog to the initial contribution PR. Updates to partner collections on GitHub are detected by hash comparison and incorporated automatically.

### Resource testing infrastructure

In the bioimage.io collection repository, GitHub Actions workflows are used to test the resources. Tests include validation of the RDF (see Resource Description File in the Online Methods) and – for models – recreation of test outputs in dynamically created test environments. As alluded to in “Contributing to the Bioimage Model Zoo”, these test results also support the maintainer’s decision to accept or block a newly submitted resource from Zenodo within a generated contribution PR.

However, these validation tests are also run regularly on already accepted resources, *e.g.,* upon release of a new bioimageio.spec or bioimageio.core version (see Developer tools in the Online Methods). This ensures long-term compatibility of the resources in the bioimage.io collection of our software tools.

The community partners can provide additional test summaries for each resource. To this end, a dedicated GitHub Actions workflow in the partner repository is triggered by the GitHub Actions workflow updating the collection. The bioimage.io bot (https://github.com/bioimageiobot) needs to be invited as a collaborator for this additional functionality. The bot can then trigger the partner’s workflow with a payload containing new or updated resource IDs from within the workflow, updating the collection in the central collection repository. The community partner is in full control over how these new or updated resources are tested. To report the generated test summaries in the bioimage.io website, they need to be included in the central test summary. Any test summary is required to include a test name and result status, as well as an error message and its traceback or warning messages as appropriate. The folder where these partner test summaries are deployed is registered in the bioimage.io collection repository, which enables each update to incorporate the latest partner test summaries and therefore, the bioimage.io website to display them in the expanded model card.

### Developer tools: bioimage.io libraries for working with models

#### Python

We provide two Python libraries for conveniently interacting with models. The first is bioimageio.spec (https://bioimage.io/docs/#/bioimageio_spec) that defines the different RDFs (see “Resource Description File”) using marshmallow (https://github.com/marshmallow-code/marshmallow). It implements programmatic access to the configuration stored in the model RDF by representing it as a Python dataclass. This representation can be created from a URL, Zenodo DOI, BioImage.IO ID or a BioImage.IO animal nickname. A crucial part of the bioimageio.spec library is its “validate” command, checking if models or other resources adhere to the RDF specification. It generates error and warning reports accordingly, which facilitates the creation of high quality metadata. Other commands include automatic upgrade of an RDF (partially) or update RDFs with another (partial) RDF. The validation, update and packaging functionalities are also available through the BioImage.IO command line tool “bioimageio”.

The bioimageio.spec library only requires minimal dependencies, which makes it easy to include it in Python applications or even Python in the browser applications, such as the model upload functionality (see Contributing to the BioImage Model Zoo in the Online Methods), which uses the bioimageio.spec library via Pyodide (https://pyodide.org/en/stable/).

The second, more advanced library is our bioimageio.core library (https://github.com/bioimage-io/core-bioimage-io-python) which requires further dependencies to implement more nuanced interactions with models. It primarily provides functionality to run inference with a BioImage.IO model for all supported weight formats except tensorflow-js. Prediction can be run for a single input batch, with padding to support input data that does not fit the model’s input shape requirements, or with tiling to support input data that is too large given the available computational resources. The standardized pre- and post-processing specified in the model RDF are automatically applied. Based on this prediction functionality, bioimageio.core also implements a test function that checks whether the expected test output can be reproduced from the test input of the associated model. Furthermore, it offers functionality to create a model RDF, package associated model data in a ready-to-use ZIP archive, and convert between selected subsets of weight formats *(e.g.,* converting tensorflow to keras weights or pytorch state dict to torchscript weights). It is implemented to be used without any deep learning framework, or only a subset of them, being installed. For example, some core functionality, like download of models, is supported without any deep learning framework installed, while prediction always depends on a deep learning framework. If only Pytorch is installed, then prediction is only possible for models containing Pytorch state dict or Torchscript weights; if Tensorflow is available as well, then also, then models with Tensorflow saved model bundle weights are supported, and so on Prediction and model test functionality are also available as command line tools. This library should be used by Python based consumer software to implement BioImage Model Zoo support. It is already used for this purpose by ilastik, the StarDist python library, ZeroCostDL4Mic and several BioEngine Apps (see “BioEngine”).

#### Java

Two general purpose Java libraries do currently exist, one lightweight core library and a library that aims to provide all required functionality to integrate with our bioimage.io infrastructure.

The lightweight library *core-bioimage-io-java* (https://github.com/bioimage-io/core-bioimage-io-java) allows model consumers and producers to load (save) to (from) the latest model RDF specification. Exported files are fit for direct upload to the bioimage.io model zoo website. The only dependency of this library is SnakeYAML (https://github.com/snakeyaml/snakeyaml), preventing any unnecessary bloating of dependencies or licensing issues for existing projects that decide to use *core-bioimage-io-java*.

The second library, *imagej-modelzoo* (https://github.com/imagej/imagej-modelzoo), builds on top of the aforementioned core library and is part of the ImageJ2^53^ and Fiji^22^ ecosystem. It is available as a Java library, but is also available on an ImageJ2/Fiji update site *(CSBDeep).* It provides the means to train and run inference of Tensorflow 1.x models. The library does this by transforming ImgLib2^54^ images into an adequate tensor representation. It also back-transforms resulting tensors into the required ImgLib2 data structures. Additionally, after installing the mentioned update site, this library’s functionality can be called directly from ImageJ macros.

When used to run model inference, *imagej-modelzoo* takes an open image in ImageJ2/Fiji and applies the standardized pre- and post-processings, as specified in the model RDF, before executing the Tensorflow 1.x compatible model as described above. The resulting output image can then be saved in any format supported by ImageJ2/Fiji.

At the time of publication, two demonstrator deep learning models are available: Noise2Void^6^ for denoising 2D or 3D images, and DenoiSeg^55^ for joint denoising and segmentation. Both demonstrator plugins can be used as blueprints for developers interested in offering similar functionality.

### BioEngine

The BioEngine application framework enables the deployment of deep learning models in the browser. It is built on top of the ImJoy plugin framework which allows connecting plugins that run across languages or servers. While BioEngine apps can run completely in the browser by using in-browser deep learning frameworks such as Tensorflow.js or ONNX.js, a typical BioEngine app for model testing consists of the web application plugin for providing user interface and the computational backend for performing the actual model inference. To enable the execution of most models in the mainstream deep learning frameworks such as Tensorflow, Keras and PyTorch, we built a BioEngine backend that uses a set of container-based server components managed within a Kubernetes cluster. The cluster is managed by a recently developed data management and AI model serving software named Hypha. We developed a custom model runner for the BioEngine using the bioimageio.core python library: the user sends requests from the BioEngine app, the requests are routed by Hypha and Nvidia Triton Inference server, and finally, processed by the model runner. The BioEngine is not only designed to be accessible from the web applications within the BioImage Model Zoo, but currently can also be accessed by other desktop software including ilastik, Icy, or QuPath. We provide the corresponding client library (https://pypi.org/project/pyotritonclient/) in Python and Javascript for performing server-side model inference.

The BioEngine can also be used to deliver more advanced functionality as web applications linked to models in the Zoo. For example, we used it to implement an application that delivers the best performing approach from a recent Kaggle challenge for protein classification in cellular images, see Use Case 3 in “Use-cases” for details. We also used it to implement the Mutex Watershed application, which provides online instance segmentation for models that predict pixel affinities. Pixel affinities correspond to a directed boundary prediction (is there a label transition across a fixed direction and pixel offset in the image) and the Mutex Watershed can turn these into an instance segmentation directly. Here, we use a version of this algorithm compiled to WebAssembly with Emscripten (https://emscripten.org/), which allows us to run it in the browser directly.

### Use-cases

**Case 1:** The StarDist model for nucleus segmentation in H&E images was trained with the StarDist python library (https://github.com/stardist/stardist) using the two published datasets^38,39^. It was then exported as model RDF using the dedicated function in the StarDist python library (https://github.com/stardist/stardist/blob/master/stardist/bioimageio_utils.py), which uses the bioimageio.core package internally (see “Developer tools: Bioimage.io libraries for working with models”). An example of this training can also be seen in the notebook at https://github.com/stardist/stardist/blob/dbd4641b78e83c37b970d2064b8bf0d5a40951a7/examples/other2D/bioimageio.ipynb. The model predicts the intermediate StarDist representations, foreground probabilities and distances, to which a non-maximum-suppression algorithm can be applied to obtain an instance segmentation. In deepImageJ, the implementation of this algorithm from the StarDist Fiji plugin is used and is invoked via an ImageJ macro. QuPath implements its own version of this algorithm and the model inference is implemented using the tensorflow java library. ZeroCostDL4Mic uses the StarDist python library internally, but offers a notebook (https://colab.research.google.com/github/HenriquesLab/ZeroCostDL4Mic/blob/master/Colab_notebooks/BioImage.io%20notebooks/StarDist_2D_ZeroCostDL4Mic_BioImageModelZoo_export.ipynb) that can be executed in Google Colab (which offers free GPU access without further configuration) without knowledge of the underlying code. The StarDist python library is the reference StarDist implementation and also contains the original implementation of the StarDist non-maximum suppression.

**Case 2:** The 3D U-Net was trained to segment cell boundaries using the 3D U-Net ZeroCostDL4Mic notebook (https://colab.research.google.com/github/HenriquesLab/ZeroCostDL4Mic/blob/master/Colab_notebooks/BioImage.io%20notebooks/U-Net_3D_ZeroCostDL4Mic_BioImageModelZoo_export.ipynb) and data from Wolny *et al*^4^. It can be directly applied to the new data from https://osf.io/fzr56/ in the ilastik neural network workflow. The network predictions are then saved and, together with the raw data, used as input to the ilastik multicut workflow. This workflow implements graph based instance segmentation and contains an interactive edge classifier. We use it to label a few edges and thus obtain a curated instance segmentation. We then use the same ZeroCostDL4Mic notebook to fine-tune the model on the data, using boundaries derived from the curated instance segmentation as target. The fine-tuned model can be exported from the notebook as model RDF, using bioimageio.core internally, and we upload it to the Zoo. The new model is applied directly in deepImageJ and its boundary predictions are post-processed to obtain an instance segmentation using the Morphological Segmentation plugin of MorphoLibJ^42^. This segmentation is used to derive instance based statistics, like the sphericity histogram.

**Case 3:** The solution is produced by the winning team *(bestfitting)* of the *Human Protein Atlas – Single Cell Classification competition* hosted on the Kaggle platform (https://www.kaggle.com/c/hpa-single-cell-image-classification/). The workflow consists of three models, one for nuclei segmentation, one for cell segmentation, and the other one for multi-label classification. The nuclei segmentation model and the cell segmentation one share the same model architecture (DPN-Unet) produced by the data science bowl 2018 nuclei segmentation competition^13^, but trained on different datasets. The top-winning teams trained the nuclei segmentation model with the data from the same competition, and the cell segmentation was trained with the HPA cell segmentation dataset^56^. In the demonstrated workflow, we first load an image from the human protein atlas (https://www.proteinatlas.org/), the nuclei channels are segmented with the nuclei segmentation model. Then, the image is fed into the cell segmentation model, the cell instances are extracted by applying a watershed-based processing (with the nuclei mask as seed). Each cell is then cropped, background and other cells are masked, and fed into the Inception-v3 based multi-label classification model. The model inference is either run locally via bioimage.core Python library running in a conda environment, or through the BioEninge backend running in our cloud computing cluster in the web application. The results are displayed with napari^32^ for the local demo and Kaibu^49^ with the remote.

**Case 4:** The mitochondria domain adaptation model is trained using torch-em (https://github.com/constantinpape/torch-em), a pytorch based library implementing deep learning approaches for microscopy image analysis. It was trained on 2D EM images of mitochondria from the Segmented anisotropic ssTEM dataset of neural tissue^57^. This model is trained to improve mitochondria foreground predictions by exploiting predictions of a shallow classifier. The model only gets these predictions as input and can thus be used in a domain adaptation setting. For mitochondria segmentation in images from a different EM modality or target tissue the user only needs to train a shallow model, which is fast and convenient due to established tools that provide interactive training functionality. Here, we demonstrate this approach with the trainable Weka^47^ plugin, Labkit^46^ and ilastik pixel classification for data from the MitoEM challenge^45^. The predictions can then be improved by applying the domain adaptation model, using either deepImageJ or the ilastik neural network workflow (any other software that supports running bioimage.io models would also work).

